# The D614G mutation in the SARS-CoV2 Spike protein increases infectivity in an ACE2 receptor dependent manner

**DOI:** 10.1101/2020.07.21.214932

**Authors:** Junko Ogawa, Wei Zhu, Nina Tonnu, Oded Singer, Tony Hunter, Amy L Ryan (Firth), Gerald M Pao

## Abstract

The SARS-CoV2 coronavirus responsible for the current COVID19 pandemic has been reported to have a relatively low mutation rate. Nevertheless, a few prevalent variants have arisen that give the appearance of undergoing positive selection as they are becoming increasingly widespread over time. Most prominent among these is the D614G amino acid substitution in the SARS-CoV2 Spike protein, which mediates viral entry. The D614G substitution, however, is in linkage disequilibrium with the ORF1b P314L mutation where both mutations almost invariably co-occur, making functional inferences problematic. In addition, the possibility of repeated new introductions of the mutant strain does not allow one to distinguish between a founder effect and an intrinsic genetic property of the virus. Here, we synthesized and expressed the WT and D614G variant SARS-Cov2 Spike protein, and report that using a SARS-CoV2 Spike protein pseudotyped lentiviral vector we observe that the D614G variant Spike has >1/2 log_10_ increased infectivity in human cells expressing the human ACE2 protein as the viral receptor. The increased binding/fusion activity of the D614G Spike protein was corroborated in a cell fusion assay using Spike and ACE2 proteins expressed in different cells. These results are consistent with the possibility that the Spike D614G mutant increases the infectivity of SARS-CoV2.

## Introduction

The SARS-CoV2 Coronavirus that initiated the current global pandemic, appeared in late 2019 in Hubei province, China(1)(2). The epidemic experienced a fast growth early on in the city of Wuhan as its epicenter in late 2019 and early January 2020 and then declined in China as a whole by the second half of February 2020. By the time the epidemic reached Europe, a variant strain had appeared that carried a missense mutation in the Spike glycoprotein that substituted the aspartate at position 614 for a glycine in isolates identified in Germany, Italy and Mexico (3). This mutation is in linkage disequilibrium with the ORF1b gene P314L substitution. In almost all cases ORF1b P314L and Spike D614G variants co-occur. The Spike glycoprotein is a type I membrane protein and the largest surface protein of the SARS-CoV2 coronavirus. It mediates infection of target cells through binding to its cognate receptor angiotensin converting enzyme 2 (ACE2) and then initiating viral-host cell membrane fusion (4). After the appearance of the Spike D614G variant in the latter course of the Chinese epidemic, over time in most examined local epidemics an enrichment of the 614G Spike protein variant over the original 614D variant has been observed, leading to the hypothesis that the Spike D614G mutation is positively selected (5)(Supplemental movie, https://nextstrain.org/ncov/global?c=gt-S_614)(6). The caveat, however, is that due to possible multiple introductions and reintroduction events a founder effect could also explain the observed viral strain dynamics. Here, we present evidence that the D614G Spike mutant displays a slightly increased infectivity (∼5X) in ACE2-expressing cells without a contribution of the ORF1b P314L mutation, when tested in pseudotyped lentiviral vectors. This result provides a plausible mechanism for the increased observed infectivity inferred from epidemiological observations and is consistent with the positive selection hypothesis of the D614G mutation.

## Results

The Spike protein is the largest structural protein of the SARS-CoV2 virus (2). This type I trimeric membrane protein mediates the viral entry into target cells through the binding of its primary receptor ACE2, and possibly also interaction with additional receptors or co-receptors (7). Activation of the Spike protein for receptor binding, requires proteolytic processing by serine proteases, such as furin, at the polybasic site during its secretion from the producer cells, or alternatively by the target cell plasma membrane TMPRSS2 protease or endosomal proteases such as cathepsins, into the S1 and S2 domains (4, 8). Upon binding to the ACE2 receptor, the S1 domain is shed and the S2 domain is exposed. The S2 domain has to be further proteolytically activated to expose the fusion peptide, which initiates membrane fusion of the viral and host cell membranes to mediate viral entry into the cytoplasm (8). The D614G mutation on the Spike protein is located at the C-terminus of the S1 fragment and outside of the receptor-binding domain, and thus is unlikely to directly influence ACE2 binding.

### Cell biology of Spike-mediated fusion

Expression of the Spike glycoprotein (GP) in 293T cells invariably led to some amount of cell fusion. This was observed in transient transfection (Figure 1A) as well as in a stable cell line generated by infection with a lentiviral vector expressing Spike GP (Figure 1B). The majority of expressed Spike GP was located in the endoplasmic reticulum (ER) as seen by calnexin colocalization in methanol-permeabilized cells (Figure 1A). Spike was also observed on the nuclear membrane and to a lesser extent on the plasma membrane. Staining cells without permeabilization showed that Spike is readily expressed on the plasma membrane, in the absence of any other viral protein, and therefore should be capable of mediating cell fusion (Fig. 1A right panel). We did not observe any significant localization differences between Spike WT and Spike D614G in transiently transfected cells. Higher levels of expression led to aberrant ER morphology probably due to ER-ER fusion events. In addition, we observed fusion of the outer nuclear membrane leading to syncytia with aggregated nuclei (Figure 1B). To test the activity of the SARS Cov2 protein in vitro, we used a cell fusion assay in which 293T cells were either transfected with Spike or transfected with the Spike receptor ACE2 (8). In order to distinguish and visualize the Spike and ACE2-expressing cells, the Spike protein was co-transfected with EGFP and the ACE2 protein cotransfected with TdTomato. Cells were mixed, plated and then observed by immunofluorescence microscopy or Fluorescence Aided Cell Sorting (FACS) 24 hours after plating. The experiments were performed with both the original Spike 614D variant GP as well as the Spike 614G mutant GP, and mouse ACE2 and human ACE2 were compared. Mixing of both mouse and human ACE2-expressing cells with Spike GP expressing cells led to an increase in syncytia frequency as well as size, i.e. syncytia with more nuclei, as observed by microscopy (Figure 1C). Both of these effects appeared to be further increased using human ACE2 compared to mouse ACE2. FACS analysis showed that transfection of Spike GP increased red/green double positive events indicative of fusion 7-8X. Inclusion of the mouse ACE2 in the TdTomato cells increased the fusions to ∼17-18X over the baseline without Spike (Figure 1E). Use of the human ACE2 only increased measured fusion events 5-6X due to the loss of the large syncytia (Figure 1E). In order to quantify the extent of fusion, we measured the depletion of syncytia rather than counting fused cells, as Spike-mediated fusion produced syncytia that were frequently large enough to be lost during passage through the FACS preparation mesh filters designed to eliminate cellular aggregates. Using this assay we observed an increase in fusion cell depletion when target cells were transfected with either mouse or human ACE2. Fusion efficiency was higher with human ACE2, but mouse ACE2 enhancement of fusion was clearly detectable by this assay, consistent with microscopy observations. The difference between Spike GP 614D and 614G with mouse ACE2 was not significant (p = 0.0722). With the human ACE2 however, we observed an 11% enhancement of D614G GFP+ cell depletion compared to WT Spike, which although small in magnitude achieved statistical significance (p <0.02) (Figure 1F).

**Figure 1.**
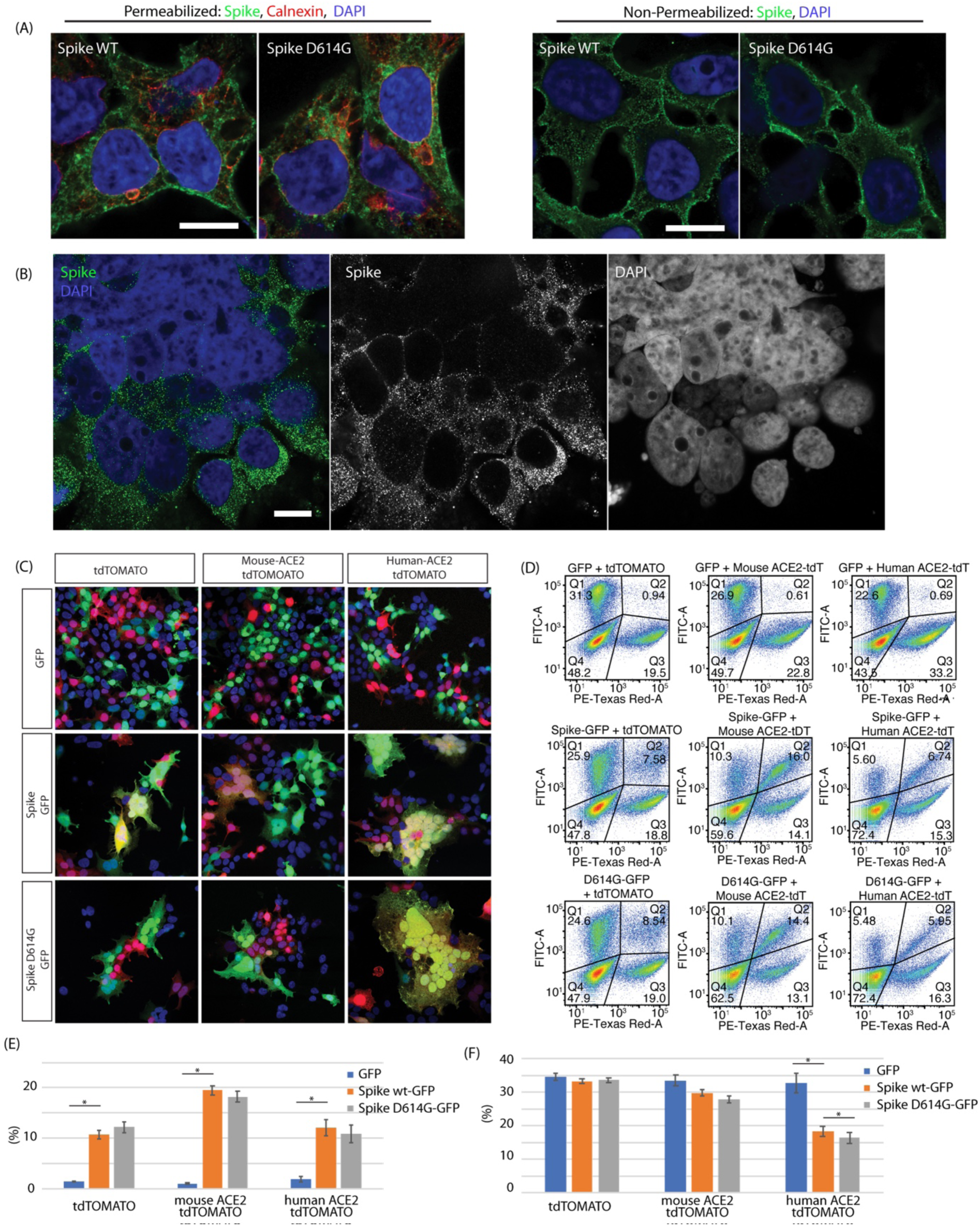
Spike protein mediated membrane fusion. (A) Immunostaining of methanol permeabilized 293T cells expressing the SARS-CoV2 Spike protein showed localization mostly in the ER (calnexin marker), but significant amounts on the plasma membrane. Non-permeabilized PFA fixed preparations showed significant amounts of Spike protein on the plasma membrane. (B) Spike protein mediated plasma membrane fusion and syncytia formation and ER-ER mediated nuclear membrane fusion in cells co-transfected with hACE2 and SARS-CoV2 Spike. (C) SARS-CoV2 Spike cell fusion assay. 293T cells were transfected with 1. Spike with EGFP or 2. ACE2 and TdTomato. Cells were then mixed and plated. Cotransfection of ACE2 increased the number and size of fused syncytia. Human ACE2 greatly increased cell fusion and hence very large syncytia. (D) FACS analysis of cell fusion experiments. Observe the depletion of the unfused Spike (GFP/FITC-A channel) population as well as the double positive (Spike GFP/ ACE2 TdTomato) population due to the formation of large syncytia. (E) FACS quantification of observed GFP+ Tdt + double positive population. Introduction of Spike and ACE2 increased fusion. As a result of increased fusion efficiency with human ACE2, depletion of the double positive population was observed due to formation of large syncytia as seen in 1C. Fusion statistics: RFP+GFP+ 293T GFP vs Spike WT-GFP p = 0.01035, mouse ACE2 GFP vs Spike WT-GFP p = 0.00135, human ACE2 GFP vs Spike WT-GFP p = 0.01604. (F) Quantification of FACS double positive (fused cells) depletion due to large syncytia loss. Without supplementation of ACE2 there was no loss of fused cells (small syncytia). With mouse ACE2 the difference between Spike 614D and 614G was not significant (p = 0.0722), whereas with human ACE2 the increased 614G syncytial depletion (−11%) as a measure of fusion achieved statistical significance (p = 0.0185). GFP depletion statistical significance: Human ACE2 GFP vs Spike WT-GFP p = 0.013232, Human ACE2 Spike WT-GFP vs Spike D614G-GFP p = 0.01879.

### Spike pseudotyped lentiviral vectors

In order to obtain a better dynamic range assay, we generated SARS-CoV2 pseudotyped lentiviral vectors that incorporated Spike GP as their envelope protein carrying a payload expressing EGFP under the CAG promoter. This pseudotyped viral vector was used to infect mouse and human ACE2 expressing cells (9). The Spike pseudotyped lentiviral vector physical titer was normalized to HIV p24 Gag protein prior to infection for all viral vectors generated. Comparison of infection efficiencies of control 293T cells with stable cells expressing either mouse or human ACE2 demonstrated an increased infection efficiency (15X) in the human ACE2 expressing cells over the 293T cell background. A similar increase was not observed in mouse ACE2-expressing cells. Comparison of Spike 614G with Spike 614D showed a consistently enhanced infectivity of the Spike 614G variant (∼5X) (Figure 2A) Co-transfection of TMPRSS2 in 293T cells did not enhance infectivity suggesting that proteolytic processing is not limiting in these cells (data not shown). Comparison of proteolytic processing of 614D and 614G Spike proteins by immunoblotting with an antibody against a juxtamembrane epitope on partially purified pseudotyped lentiviral vector virions showed neither enrichment of Spike incorporation nor proteolytic processing differences. This was observed both with partially purified viral particles as well as in viral supernatants (Figure 2B).

**Figure 2.**
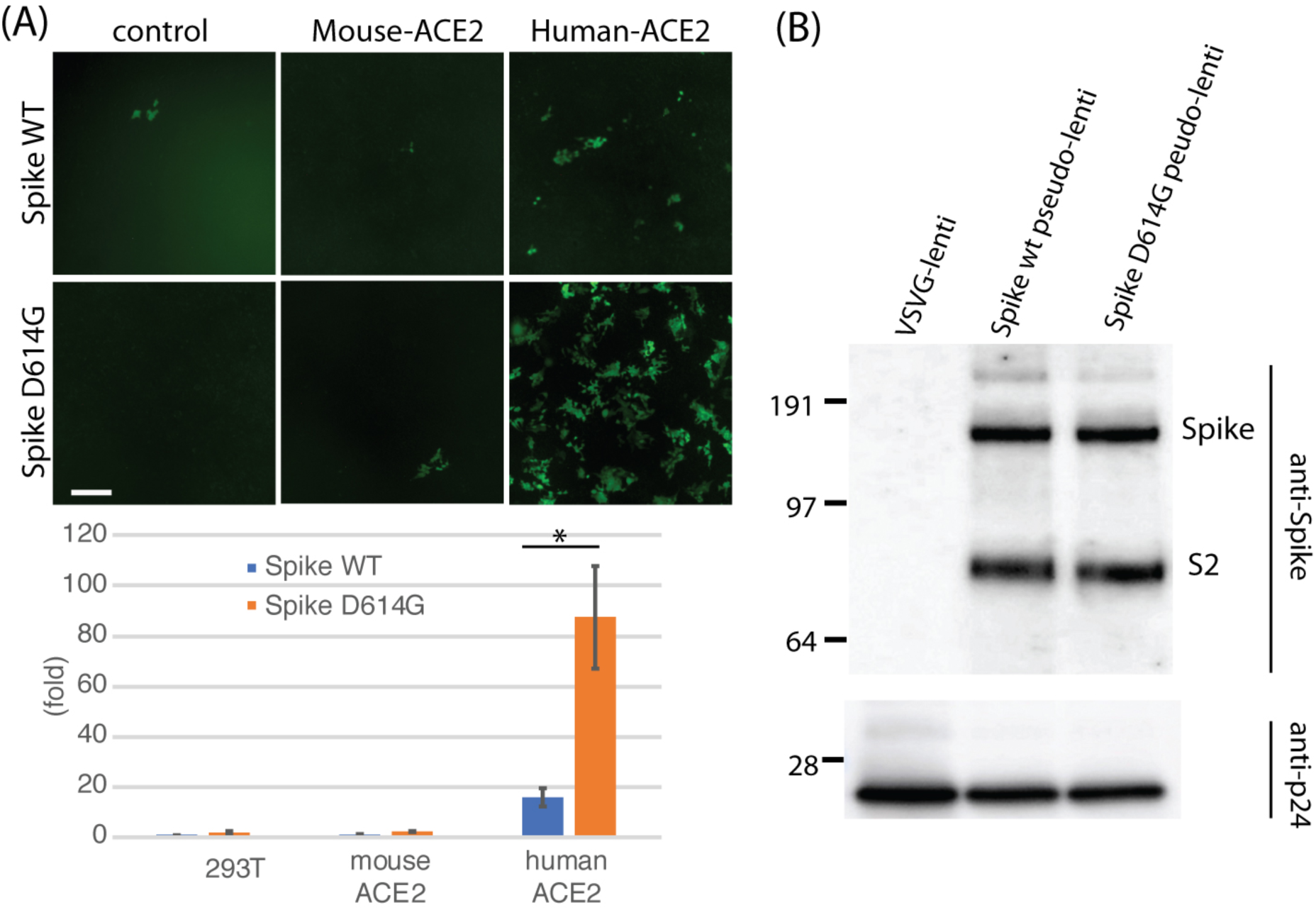
D614G infectivity enhancement in pseudotyped lentiviral vectors. (A) Lentiviral vector pBOB-CAG-GFP was pseudotyped with either WT Spike or D614G Spike, titered and normalized by the level of HIV p24 Gag protein and used to infect control 293T, and 293T cells stably expressing mouse ACE2 or human ACE2. Fluorescence microscopy of GFP (top) and FACS quantification (bottom) are shown. The D614G mutation increased infectivity of Spike ∼5X (p = 0.0471) by FACS. Baseline infection of 293T control cells with pseudotyped 614D spike pseudotyped lentiviral vector is set as 1X (B) Immunoblot of partially purified lentiviral preparations shows that incorporation of Spike protein did not appreciably change with the D614G mutation, nor was proteolytic processing of the S2 fragment different. Viral vector nucleocapsid was quantified using a monoclonal antibody against the HIV p24 Gag protein.

## Discussion

The emergence of the SARS-CoV2 Spike D614G and its rapid increase in prevalence has been striking (5). Observational population genetics alone could not resolve whether this mutation and/or the frequently co-occurring ORF1b P314L allele alter biological activity or are just an epiphenomenon underlying a founder effect due to repeated introductions. Here, in the present report we find functional evidence that the Spike D614G mutant has modestly different biochemical properties consistent with positive selection of the D614G mutant SARS-CoV2 virus variant. The increase in cell entry activity of pseudotyped lentiviral vectors and cell fusion allowed us to dissect the Spike protein function in isolation and test the activity of the D614G point mutation. In our work we observed no increases in expression, increased stoichiometry of Spike incorporation into pseudotyped viral vector particles, nor proteolytic processing that could account for the increased infectivity. In contrast, other reports have respectively reported increases and decreases in S2 proteolytic processing (10–14). Given the location of the 614 residue within the S1 fragment distal to the RBD, it is also unlikely that the D614G mutation directly affects the ACE2 binding activity. Also unlikely is an affect on the S2’ proteolytic processing that activates the fusion peptide, as the S1 fragment with the 614 site is already released before S2’ proteolytic processing can occur. Not yet investigated are the post-binding steps leading to the release of S1 before the S2’ activation of the fusion peptide. Thus, a biochemical basis for increased infectivity is observed in the SARS-CoV2 D614G mutant consistent with positive selection observed in the field. Nevertheless, we cannot currently explain the basis for this increased infectivity, and therefore further work is needed to clarify the mechanistic basis of the increased infectivity.

## Methods

Pseudotyped lentiviral vectors were prepared, titered and partially purified as described in Tiscornia et al. (15). For small scale production, calcium phosphate transfection was substituted for Lipofectamine LTX using the manufacturer’s recommended conditions. The Spike protein gene sequence was synthesized based on the original US isolate SARS-CoV2 hCoV19_USA EPI_ISL_414366 (GISAID) with human codon optimization and the addition of a C-terminal HA tag (IDT), and cloned by Gibson assembly using the NEB kit into the lentiviral pBOB-CAG vector. The D614G substitution was introduced with a single gBlock spanning the AspI and RsrII sites and Gibson assembly (NEB). Pseudotyping plasmids were generated by deletion of the external CMV promoter and 5’ LTR leaving the internal CAG promoter by restriction digestion, and self-ligation.

A stable Spike expressing 293T cell line was generated by infection with pBOB-CAG-SARS-CoV2-Spike-HA lentiviral vector https://www.addgene.org/141347/ followed by single cell cloning and screening by cell extract immunoblotting of individual clones with HA tag antibody (CST #3724). Immunofluorescence was performed following the Cell Signaling Technology protocol (https://www.cellsignal.com/). Primary antibodies were GeneTex 632604 anti-Spike antibody (mouse monoclonal) and anti-calnexin (C5C9) rabbit mAb (CST #2679) were detected with 2^nd^ anti-Mouse IgG Alexa 488 (Life tech Cat# A11029) and anti-rabbit IgG Dylight 633 (Thermo Fisher #35563) antibodies. Imaging was performed with Zeiss AiryScan 880 microscope. Human and mouse ACE2 expressing cell lines were generated by lentiviral vector (pLV-human-ACE2 IRES Puro or pLV-mouse-ACE2 IRES Puro, gifts from Dr. Sumit Chanda) transduction followed by puromycin selection. Statistical analyses were performed using Fisher’s 2 tailed T-test on MS Excel.

## Author Contributions

GMP conceived the project. JO, & GMP designed experiments. JO and GMP performed experiments with technical support of NUT. WZ and GMP generated the Spike expressing stable cell line. OS proposed the fusion assay. ALR conceived and performed the lung epithelial experiments. TH provided scientific advice and logistic support. GMP, JO, TH and ALR wrote the manuscript.

## Acknowledgements

This work was supported by a generous donation from Danielle A Fong. JO, NUT and GMP are supported by a medical research grant from the WM Keck foundation. OS is supported by the (2016/ Weizmann Institute Staff Scientists Internal Grant Program) ALR is supported by the Hastings Foundation. TH is an American Cancer Society Professor and holds the Renato Dulbecco Chair. Funding from NCI CA014195 is acknowledged for core support.

## References

1. P. Zhou, et al., A pneumonia outbreak associated with a new coronavirus of probable bat origin. Nature 579, 270–273 (2020).

2. F. Wu, et al., A new coronavirus associated with human respiratory disease in China. Nature 579, 265–269 (2020).

3. P. Stefanelli, et al., Whole genome and phylogenetic analysis of two SARSCoV-2 strains isolated in Italy in January and February 2020: Additional clues on multiple introductions and further circulation in Europe. Eurosurveillance 25, 1–5 (2020).

4. M. Hoffmann, et al., SARS-CoV-2 Cell Entry Depends on ACE2 and TMPRSS2 and Is Blocked by a Clinically Proven Protease Inhibitor. Cell 181, 271-280.e8 (2020).

5. B. Korber, et al., Spike mutation pipeline reveals the emergence of a more transmissible form of SARS-CoV-2. bioRxiv 4, 2020.04.29.069054 (2020).

6. J. Hadfield, et al., NextStrain: Real-time tracking of pathogen evolution. Bioinformatics 34, 4121–4123 (2018).

7. J. L. Daly, et al., Neuropilin-1 is a host factor for SARS-CoV-2 infection. bioRxiv, 2020.06.05.134114 (2020).

8. S. Belouzard, V. C. Chu, G. R. Whittaker, Activation of the SARS coronavirus spike protein via sequential proteolytic cleavage at two distinct sites. Proc. Natl. Acad. Sci. U. S. A. 106, 5871–5876 (2009).

9. X. Ou, et al., Characterization of spike glycoprotein of SARS-CoV-2 on virus entry and its immune cross-reactivity with SARS-CoV. Nat. Commun. 11 (2020).

10. L. Zhang, et al., The D614G mutation in the SARS-CoV-2 spike protein reduces S1 shedding and increases infectivity https://doi.org/10.1101/2020.06.12.148726.

11. Z. Daniloski, X. Guo, N. E. Sanjana, The D614G mutation in SARS-CoV-2 Spike increases transduction of multiple human cell types https://doi.org/10.1101/2020.06.14.151357.

12. J. Hu, et al., The D614G mutation of SARS-CoV-2 spike protein enhances viral infectivity 1 and decreases neutralization sensitivity to individual convalescent sera 2 Running Title: D614G mutant spike increases SARS-CoV-2 infectivity https://doi.org/10.1101/2020.06.20.161323.

13. S. Ozono, et al., Naturally mutated spike proteins of SARS-CoV-2 variants show differential levels of cell entry. bioRxiv, 2020.06.15.151779 (2020). https://doi.org/10.1101/2020.06.15.151779

14. S. Isabel, et al., Evolutionary and structural analyses of SARS-CoV-2 D614G 3 spike protein mutation now documented worldwide https://doi.org/10.1101/2020.06.08.140459.

15. G. Tiscornia, O. Singer, I. M. Verma, Production and purification of lentiviral vectors. Nat. Protoc. 1, 241–245 (2006).

